# Recombination proteins differently control the acquisition of homeologous DNA during *Bacillus subtilis* natural chromosomal transformation

**DOI:** 10.1101/2020.05.13.090589

**Authors:** Ester Serrano, Cristina Ramos, Juan C. Alonso, Silvia Ayora

## Abstract

In naturally competent *Bacillus subtilis* cells the acquisition of closely related genes occurs *via* homology-directed chromosomal transformation (CT), and its frequency decreases log-linearly with increased sequence divergence (SD) up to 15%. Beyond this and up to 23% SD the interspecies boundary prevails, the CT frequency marginally decreases, and short (<10-nucleotides) segments are integrated *via* homology-facilitated micro-homologous integration. Both poorly known CT mechanisms are RecA-dependent. Here we identify the recombination proteins required for the acquisition of interspecies DNA. The absence of AddAB, RecF, RecO, RuvAB or RecU, crucial for repair-by-recombination, does not affect CT. However, inactivation of *dprA, radA, recJ, recX* or *recD2* strongly interfered with CT. Interspecies CT was abolished beyond ~8% SD in Δ*dprA*, ~10% in Δ*recJ*, Δ*radA*, Δ*recX* and 14% in Δ*recD2* cells. We propose that DprA, RecX, RadA/Sms, RecJ and RecD2 help RecA to unconstrain speciation and gene flow. These functions are ultimately responsible for generating genetic diversity and facilitate CT and gene acquisition from bacteria of the same genus.

## Introduction

The unidirectional nonsexual incorporation of DNA into the genome of a recipient bacterium (horizontal gene transfer) has been the major evolutionary force that has constantly reshaped genomes and persisted through evolution to maximize adaptation to new ecological conditions and genetic diversity ^1;Fraser, 2007 #154^. The exchange of chromosomal genes among members of the same or related species occurs *via* RecA-dependent homologous recombination (HR) by Hfr conjugation, viral transduction or natural chromosomal transformation (CT) ^2–4^. In Hfr conjugation and transduction the recombination machinery processes double-stranded (ds) DNA, whereas during CT the machinery integrates single-stranded (ss) DNA ^4,5^. Bacteria have developed mechanisms to fight gene transfer by conjugation and transduction, but cells are less efficient to impose barriers to natural transformation ^1^. Restriction-modification systems rapidly fragmentate internalised non-self dsDNA, but ssDNA, internalised by transformation, is refractory to most restriction systems ^6,7^ Adaptive immune systems, as CRISPR-Cas, are usually absent in naturally competent bacteria ^8^.

Genes devoted to natural competence are encoded in the genome of many bacteria, among them the model organism *Bacillus subtilis*, which occupies a wide range of aquatic and terrestrial niches, and colonizes animal guts ^9^. Upon stress, a small fraction of *B. subtilis* stationary phase cells induce competence. During competence development, ongoing DNA replication is halted, a transcriptional programme is activated, a DNA uptake apparatus is assembled at a cell pole, and lysis of kin is induced, releasing DNA for uptake ^7,10–12^. This DNA uptake apparatus binds any extracellular dsDNA, degrades one strand, and internalises the complementary strand into the cytosol ^10–12^.

Naturally competent cells can integrate fully homologous DNA (homogamic CT), and with lower efficiency homeologous (similar but not identical) DNA (heterogamic CT) ^2^. This interspecies CT can replace the recipient by a homeologous DNA, leading to mosaic structures ^13–15^. The transformation frequency of heterogamic DNA decreases with increased sequence divergence (SD) both in *B. subtilis* and *S.pneumoniae* cells ^14,15^. In *B. subtilis*, the transformation assays with donor DNAs from different *Bacilli* species revealed a biphasic curve. Interspecies CT decreased log-linear up to 15% SD, and integration in this part of the curve occurred *via* one-step homology-directed RecA-dependent gene replacement ^16,17^ Beyond 15% SD the CT reached a plateau, and the integration of just few nucleotides, at micro-homologous segments (<10-nucleotides [nt]), was observed at a very low efficiency (~1 x 10^-5^) ^17^, suggesting another recombination mechanism.

CT also contributes to expand the pan-genome of a species, because it allows the integration of a heterologous DNA sequence, if flanked by two regions identical with recipient, usually >400-nt, but longer homologous regions significantly increase the integration frequency ^7,18^. Here, two homologous recombination (HR) events in the flanks integrate the heterologous DNA *via* recipient-deletion /donor-insertion with ~10-fold lower efficiency than homologous gene replacement ^7,9^. Another mechanism is observed in replicating competent *Streptococcus pneumoniae* or *Acinetobacter baylyi* cells. Here, homology at only one flank (anchor region) facilitates illegitimate recombination of short (<10-nt) micro-homologous segments with subsequent deletion of the intervening host DNA and integration of long heterologous DNA segments, albeit with very low efficiency when compared to homogamic CT ^20,21^. The length of the anchor region affects the efficiency of this homology-facilitated illegitimate recombination (HFIR) ^19^.

The proteins responsible for the acquisition of natural homeologous DNA remain poorly known. The main player during HR is the RecA recombinase. RecA from a naturally competent bacterium (*e.g., B. subtilis*) has evolved to catalyse strand exchange in either the 5’→3’ or 3’→5’ direction and to tolerate 1-nt mismatch in an 8-nt region ^16,17^ In naturally competent cells, the essential SsbA and competence-specific SsbB coated the incoming linear ssDNA as soon it leaves the entry channel. *B. subtilis* RecA cannot filament onto SsbA- or SsbB-coated ssDNA ^22,23^. The RecA accessory proteins are divided into four broad classes. First, the mediators that act before RecA-mediated homology search. The DprA mediator facilitates partial disassembly of SsbA and SsbB from the ssDNA, allows RecA binding, and in concert with SsbA assists RecA to catalyse DNA strand exchange ^23^. In the absence of DprA, RecO assists RecA to filament onto SsbA-coated ssDNA ^22^. Second, the modulators, which act during DNA identity search and strand exchange, and either promote RecA nucleoprotein filament assembly as RecF (in the *DrecX* context) or its disassembly from the ssDNA, as RecX or RecU (in the *DrecX* context) ^24–28^. With the help of both, mediators and modulators, a dynamic RecA-ssDNA filament is engaged in sequence identity search. *E. coli* RecA requires ~15-nt of homologous ssDNA to promote strand exchange, defining the *in vitro* minimal efficient processing segment (MEPS) ^29,30^. *In vivo, E. coli* RecA significantly recombines two duplex DNAs with 25- to 30-bp MEPS ^31^, thus this length was proposed for the *B. subtilis* protein ^32^. *In vivo*, upon finding a MEPS, RecA initiates strand invasion to form a heteroduplex, known as displacement loop (D-loop) ^33^. Then, branch migration translocases (RecD2, RuvAB, RecG and RadA/Sms) allow D-loop extension, and help to generate a stable heteroduplex ^28,34^. Finally, after DNA strand exchange a structure-specific nuclease must resolve the D-loop. The RecU resolvase is unable to cleave D-loops ^35–37^, and the enzyme responsible for such activity remains unknown. The contribution of these RecA accessory proteins to homogamic CT has been studied in deep ^7,28,38^, but little is known about their role in the integration of heterogamic DNA. In *B. subtilis*, two exonucleases, the AddAB complex (counterpart of *E. coli* RecBCD) and RecJ are crucial to process double-strand breaks (DSBs) during DNA repair ^33,39^, but their putative role in the degradation of the displaced strand during CT remains to be documented.

DNA transfer occurs in ecologically cohesive communities. In this study we aimed to identify the recombination mechanisms and the proteins that contribute to heterogamic CT. We found that interspecies CT frequency is similar to *rec*^+^ cells in the *addAB*, *recO*, *recF*, *ruvAB* and *recU* context. Our results suggest that for interspecies CT only a subset of recombination proteins is required. RecA, RecX, RecJ, RecD2, DprA and RadA/Sms participate in both, homology-directed HR and homology-facilitated micro-homologous integration mechanisms that are active depending on the degree of SD in *B. subtilis* competent cells.

## Results

### Experimental design

The rational to select *B. subtilis* competent cells to analyse how the recombination machinery contributes to heterogamic CT is summarised in Supplementary *Annex 1*. Competence development is a stochastic process driven by the expression of ComK, and it occurs only in a small fraction (0.1 - 5%) of cells ^11,12^. Interspecies CT, at SD above 8%, is at the limit of detection in wild type cells ^14^. To overcome such technical difficulty, Rok, which directly represses *comK* expression, was deleted ^40^. Inactivation of *rok* increases the subpopulation of cells that develop natural competence ^41^, and thereby the sensitivity of the assay. The different *rec* mutations were mobilised into the *rok* strain (see Table 1). All the strains additionally lack Rok, but for simplicity we only state the *rec* gene that is mutated.

**Table 1.** Homogamic CT frequency of *rec*-deficient strains

| Strain name<sup>a</sup> | Relevant genotype<sup>a</sup> | Homogamic Transformation | source |
| --- | --- | --- | --- |
| BG1359 | $\Delta rok, rec^+$ | 100 ($3.3 \times 10^{-4}$) | 17 |
| BG1641 | + $\Delta recO$ | $76 \pm 25$ | This work |
| BG1611 | + $recF15$ | $78 \pm 17$ | This work |
| BG1631 | + $\Delta addAB$ | $49 \pm 10$ | This work |
| BG1485 | + $\Delta ruvAB$ | $58 \pm 11$ | This work |
| BG1653 | + $\Delta recU$ | $34 \pm 12$ | This work |
| BG1813 | + $\Delta recJ$ | $1.2 \pm 0.9$ | This work |
| BG1397 | + $\Delta recX$ | $0.2 \pm 0.1$ | This work |
| BG1549 | + $\Delta recD2$ | $0.1 \pm 0.05$ | This work |
| BG1647 | + $\Delta radA$ | $0.06 \pm 0.003$ | 60 |
| BG1811 | + $\Delta dprA$ | $0.004 \pm 0.002$ | This work |
| BG1633 | + $\Delta recA$ | <0.001 | 16 |

<sup>a</sup>All *B. subtilis* *rec*-deficient strains are isogenic with BG1359. The genotype of the BG1359 strain is *trpCE metA5 amyE1 ytsJ1 rsbV37 xre1 xkdA1 att<sup>SPB</sup> att<sup>ICEBs1</sup> $\Delta rok$* . This strain lacks restriction-modification, CRISPR-Cas systems, different prophages and MGEs that might reduce the transformation rate.

The *rpoB* gene encodes for the essential ß subunit of the RNA polymerase. The 2997-bp donor DNAs have a mutation at codon 482 that confers rifampicin resistance (Rif^R^, *rpoB*482) (Fig. S1B). The *rpoB*482 DNA from *B. subtilis* 168 with 1 mismatch (*Bsu* 168 *rpoB*482) was used for homogamic CT (*i.e*, the recipient’s own DNA, with just the Rif^R^ mutation). For heterogamic CT a fixed concentration of purified *rpoB* 482 donor DNAs derived from the *B. subtilis* clade (2.47% SD and 8.35% SD), the *B. amyloliquefaciens* clade with 10.12% SD, the *B. licheniformis* clade (14.52% SD and 17% SD), the *B. thuringiensis* clade with 20.83% SD, or a far distant *Bacillus* with 22.74% SD (*B. smithii* DSM4512) (Supplemental *Annex 2*, Fig. S1A). A further description of these DNAs is presented in Supplementary material *Annex 2*. As revealed in Fig. S1B, the mismatches in these natural homeologous DNAs are almost homogeneously distributed ^16,17^, although there are some short regions with higher SD (see below). To avoid fitness costs, the promoter of the *rpoB* gene is provided by the recipient strain, and the sequence identity at the protein level varied from 99% to 89% (Fig. S1C).

### DprA, RecX, RadA/Sms, RecJ and RecD2 contribute to homogamic CT, but not RecF, RecO, RecU, RuvAB and AddAB

Except in *ΔrecD2* cells, the number of Rif^R^ spontaneous mutant colonies, which appeared in the absence of *rpoB*482 DNA in the different *rec*^−^ strains was similar to the *wt* control (see Supplementary material *Annex 3*). To test how the different *rec*^-^ mutants affect intraspecies CT, competent cells were transformed with *Bsu* 168 *rpoB*482 DNA with direct selection for Rif^R^ (Table 1, Fig. 1A). The homogamic CT frequency with this donor DNA was similar to the one obtained with *Bsu* 168 *met^+^* DNA, also with a single donor-recipient mismatch ^17^. As previously reported, intraspecies CT was blocked in the *DrecA* strain (Table 1, Fig. 1A) ^16,42^.

**Figure 1.**
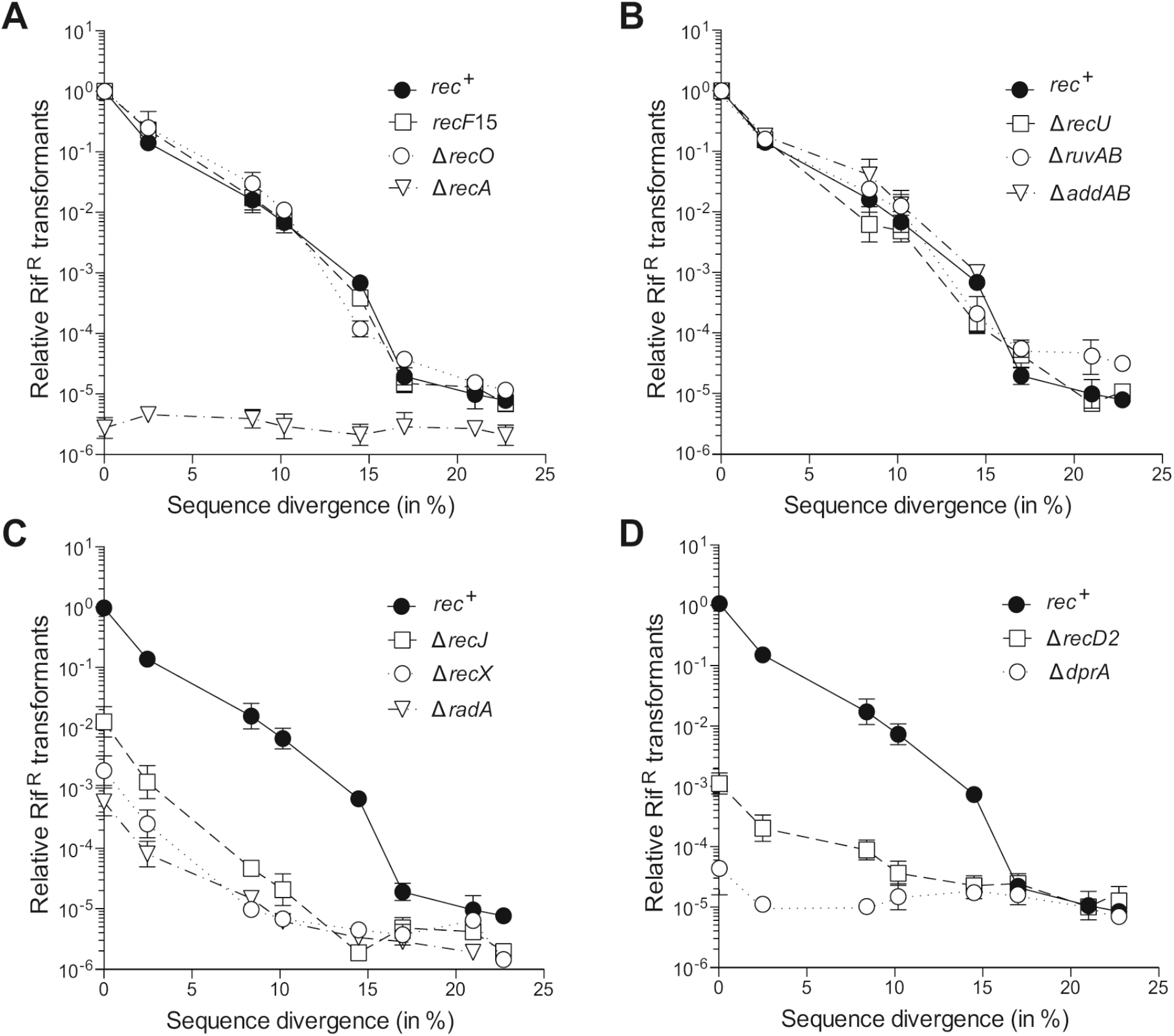
CT frequencies as a function of SD in different *rec* mutants. Donor DNA was a *rpoB*482 DNA conferring Rif; derived from *B. subtilis* 168 (0.04% SD, homologous DNA), *B. subtilis* W23 (2.47%), *B. atrophaeus* 1942 (8.35%), *B. amyloliquefaciens* DSM7 (10.12%), *B. licheniformis* DSM13 (14.52%), *B. gobiensis* FJAT-4402 (17%), *B. thuringiensis* MC28 (20.83%) and *B. smithii* DSM4216 (22.74% SD). The *rpoB482* DNA (0.1 μg DNA/ml) from these different *Bacillus* species with the selectable Rif mutation was used to transform BG1359 (*rec*^+^, ●) competent cells and its isogenic derivatives. The values are plotted dividing the number of transformants/CFUs obtained in each condition by the number of transformants/CFUs obtained when the *rec^+^* cells are transformed with *Bsu*168 *rpoB*482 DNA. In (A): BG1611 (*recF*15, □), BG1641 (*ΔrecO*,○) and BG1633 (*ΔrecA*, ▽); in (B): BG1653 (*ΔrecU*, □), BG1485 (*ΔruvAB*, ○) and BG1631 (*ΔaddAB*, ▽); in (C): BG1813 (*ΔrecJ*, □), BG1397 (*ΔrecX*, ○) and BG1647 (*DradA*, ▽); in (D): BG1549 (*ΔrecD2*, □) and BG1811 (Δ*dprA*, ○). All data points are mean ± standard error of the mean (SEM) derived from 3 to 5 independent experiments

Natural homogamic CT was barely affected in *recF15*, Δ*recO*, Δ*recU*, Δ*ruvAB*, Δ*addAB* cells (Table 1, Fig. 1A-B). Their CT frequencies were similar to results previously observed for these recombination mutants in the *rok^+^* background ^28,43^. However, the frequency of intraspecies CT was significantly reduced in competent Δ*recJ* (by ~80-fold), Δ*recX* (~500-fold), Δ*recD2* (~950-fold), Δ*radA* (~1600-fold) and Δ*dprA* (~24,000-fold) cells (Table 1, Fig. 1C-D). Except in Δ*recJ*, intraspecies CT efficiency in these mutants was lower in Δ*rok* than in *rok^+^* ^26,34^. The deleterious effect of additionally mutate *rok* in these backgrounds will be reported elsewhere.

### Interspecies CT requires DprA, RecX, RadA/Sms, RecD2 and RecJ

To gain insight into the contribution of these recombination proteins to interspecies CT, we used *rpoB*482 DNAs with different degree of SD (Supplementary *Annex 2*). The frequency of interspecies CT decreased logarithmically with increased SD up to ~15% in competent *recF*15, Δ*recO*, Δ*recU*, Δ*ruvAB* or Δ*addAB* cells (Fig. 1A-B). When SD was further increased, beyond 15% and up to ~23% SD, the interspecies CT frequencies varied <3-fold in *recF*15, Δ*recO*, Δ*recU*, *ruvAB* and *addAB* (Fig. 1A-B), as in the *rec^+^* control (Fig. 1A) ^17^, suggesting that these functions do not limit genetic recombination in otherwise *rec^+^* cells.

A different outcome was observed in Δ*recJ*, Δ*recX*, Δ*radA*, Δ*recD2* and Δ*dprA*. The CT frequency was similar to the frequency of spontaneous Rif^R^ mutations beyond ~8% SD in Δ*dprA*, ~10% in Δ*recX* and Δ*radA*, and ~15% in Δ*recJ* and Δ*recD2* competent cells, (Fig. 1C-D). We believe that this strong defect was not due to an impairment in fitness cost. The colony size observed after overnight growth at 37°C under selective pressure was similar in all cases. Furthermore, the sequence analysis of the chimeric *rpoB*482 genes revealed a RpoB482 protein only bearing the Rif^R^ mutation (see below).

### The interspecies barrier is altered in *ΔrecJ, ΔrecX, ΔrecD2, DradA* and Δ*dprA*

To simplify the analysis of the strains that are significantly impaired in heterogamic CT, we gave a value of 1 to the transformation rate obtained by the given mutant in the intraspecies CT assay, and plotted the data relative to this value. Hence only the impact of SD is evaluated (Fig. 2A-B). Different outcomes were observed. First, in competent Δ*recX*, Δ*radA* and Δ*recJ* cells interspecies CT decreased logarithmically with increased SD, but only up to ~10% (Δ*recJ*, ~14%) SD, to reach a plateau at higher divergence (Fig. 2A). Second, competent *recD2* reached a plateau already at 8% SD (Fig. 2B). Third, upon inactivation of *dprA*, the decline in the rate of recombination with SD was only observed at ~2% SD, to reach a plateau at higher divergence (Fig. 2B).

**Figure 2.**
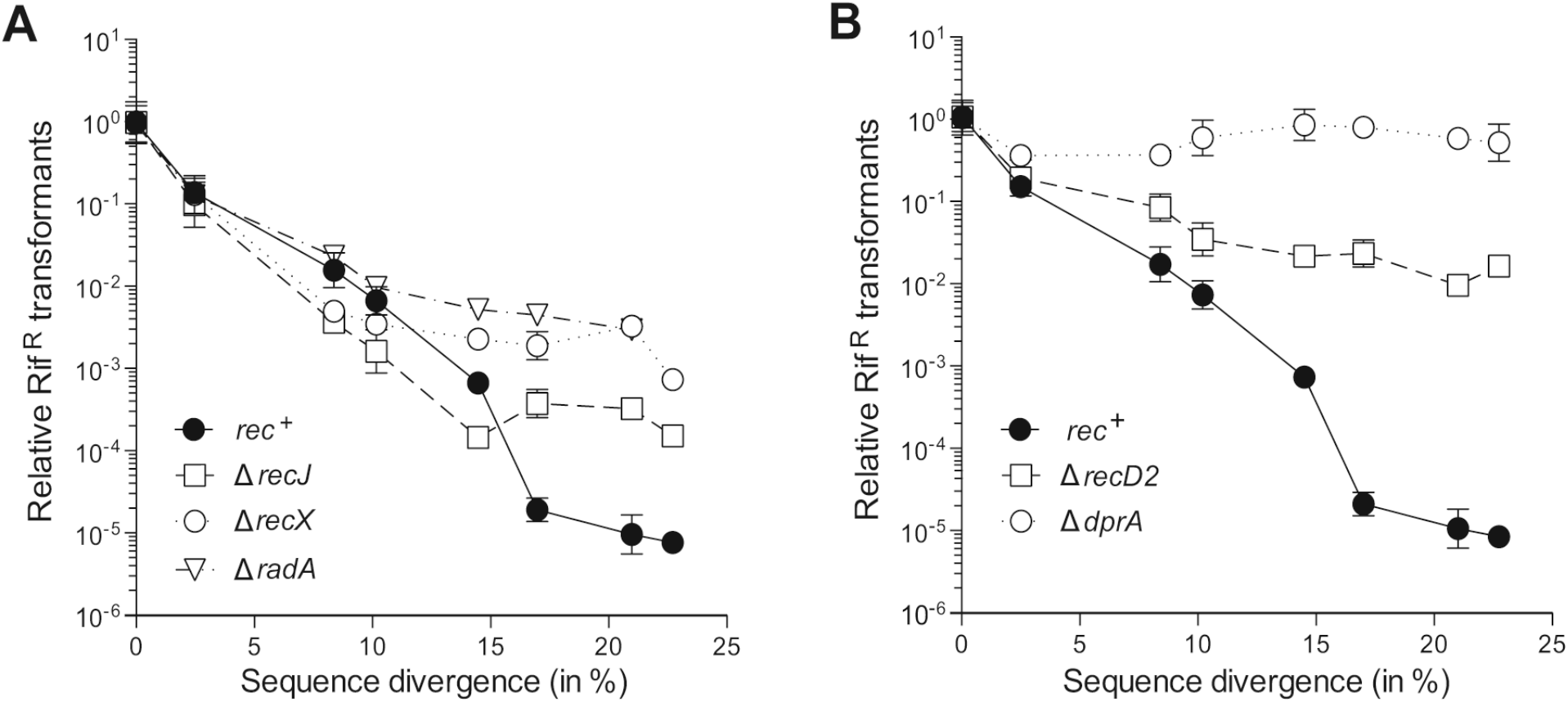
Normalised CT frequencies. The CT frequencies were normalized to give a value of 1 to the transformation frequency of the indicated *rec*-deficient strain when transformed with homogamic DNA, and heterogamic CTs are plotted relative to these values. In (A): Δ*recJ*, Δ*recX* and Δ*radA* (A); in (B): Δ*recD2* and Δ*dprA* Rif^R^ cells. For comparison, the values obtained for the *rec^+^* strain are also plotted.

### RecO, RecF, RecU, RuvAB and AddAB are dispensable in heterogamic transformation

The above results suggest that RecO, RecF, RecU, RuvAB and AddAB play no apparent roles in interspecies CT, but their contribution to the integration length is unknown. To analyse this, we sequenced the *rpoB*482 gene from Rif^R^ clones and calculated the mean integration length.

The maximal integration length that can be detected with the donor DNA of ~2% SD (*Bsu* W23 DNA) is 2628-bp, because the first mismatch is located at position 350 and the last at position 2978. Previous analysis showed that when the *rec*^+^ strain was transformed with this donor DNA, the mean integration length was close to this maximal integration length, around ~2300-bp ^16,17^. Similar results were obtained with the donor DNA of ~2% SD in the Δ*recO*, Δ*recU*, Δ*ruvAB* and Δ*addAB* backgrounds (Fig 3A-B).

**Figure 3.**
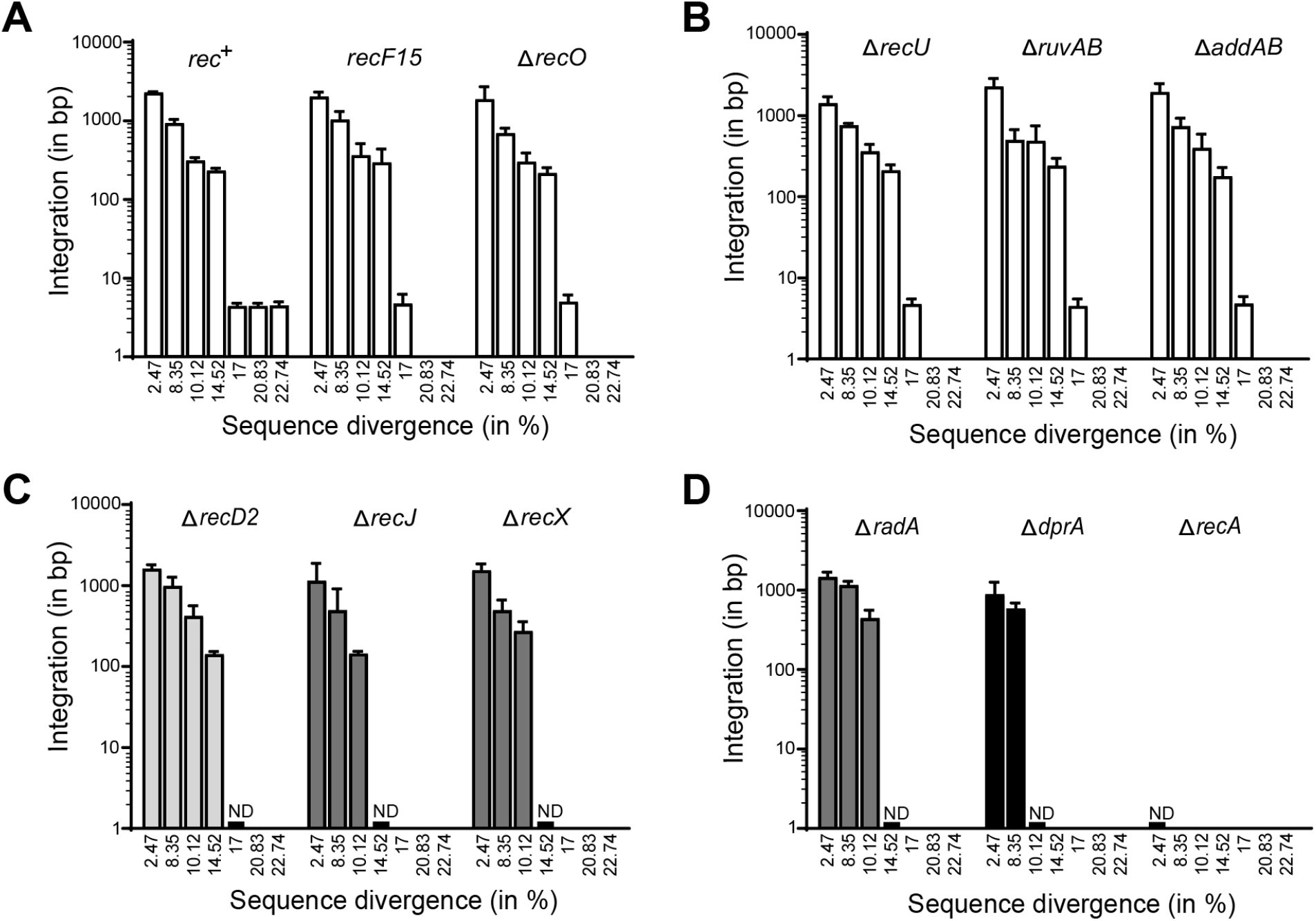
Determination of integration length of donor DNA with increased SD in different *rec* mutants. The mean length of integration was determined by sequencing the 2997-bp *rpoB*482 DNA from different transformants obtained in *rec^+^, recF15* and Δ*recO* (A); Δ*recU*, Δ*ruvAB* and Δ*addAB* (B); Δ*recD2*, Δ*recJ* and Δ*recX* (C) and Δ*radA*, Δ*dprA* and Δ*recA* (D). Integration endpoints were defined as the average between the last single nucleotide polymorphism present and the next SNP absent in the sequence of trasformants (integration endpoint). The empty bars highlight the values obtained with strains that undergo both homology-directed and homology-facilitated CT, and filled bars denote strains impaired in homology-directed and blocked in homology-facilitated CT. ND, not detected.

Up to 15% divergence range, RecA is believed to integrate the DNA by a homology-directed HR mechanism, initiating recombination in a MEPS ^16,22^. A comparison of the nucleotide sequence of donor with recipient DNA revealed the presence of 22 MEPS at or above 25-nt in donor DNA with ~8% SD, and 21 at ~10% SD. In both donor DNAs there is a long stretch of ~200-nt of sequence identity upstream of the *rpoB*482 mutation, and several regions with a MEPS longer than 54-nt downstream of the *rpoB*482 mutation. They could define the left and right recombination endpoints, being the region in-between integrated independently of its SD, as it occurs in the insertion of heterologous DNA by two-step HR at the flanks (see Introduction). However, integrated fragments of ~1600-nt (*i*.*e*, recombination endpoints at the 157-nt [at position 1-157] and the 81-nt MEPS [at position 1509-1589]) was not observed in *rec^+^* transformations with ~8% SD donor DNA, and the same was observed with 10% SD (Fig 3A). This result suggests that the two-step deletion/insertion CT may not take place with interspecies DNA, probably because it requires two longer flanking homologous regions (see Introduction), or the sequence in-between plays a relevant role.

The analysis of 10-20 Rif^R^ clones obtained in the transformation of competent *recF*15, Δ*recO*, Δ*recU*, Δ*ruvAB* and Δ*addAB* cells with donor DNA of ~8% SD revealed that the mean integration length was as in *rec*^+^, 700-900-nt, except in *ΔruvAB*, which was ~490-nt (*i*.*e*, ~2 times less) (Fig. 3A-B, and Table S1). A sequence analysis of the integrated region revealed eight MEPS (four upstream [50-, 35-, 38- and 36-nt] and four downstream [35-, 81-, 41- and 33-nt]) of the *rpoB*482 mutation. One recombination endpoint was usually at one of these MEPS, but the other endpoint was usually at other region, with a size below MEPS (Table S1). Several hypotheses can explain these results: i) the MEPS used *in vivo* are shorter, ii) RecA is insensitive to 1-nt mismatch every ~8-nt, but not to higher mismatches. Indeed, if 1-nt mismatch is allowed at the short endpoints, longer MEPS regions can be predicted in all cases, and iii) integration starts at one MEPS and proceeds uni- or bidirectionally until RecA finds a barrier (*e.g*., 2-nt mismatch every *~*8-nt). We noticed that the DNA inserts are usually followed by higher local SD (~20% to ~28%) in the next 25-nt interval, suggesting that recombination ended there because RecA found a heterologous block.

In *recF*15, Δ*recO*, Δ*recU*, Δ*ruvAB*, Δ*addAB* and *rec^+^* cells, when SD is ~10%, the mean integration length was ~300-nt (Fig. 3A-B). In these transformants, the nucleotide sequence changes do not alter the RpoB482 protein sequence (see Fig. S1C). Within a 600-nt interval centred at the *rpoB*482 mutation, there are only three sequences at or above MEPS (see Fig. 4A). Analysis of 10 different Rif^R^ clones in *rec^+^* showed that in ~70% of the cases, the 3’-endpoint was at the 54-nt MEPS (at position 1536-1590, Fig. 4A). Recombination in this region did not extend beyond, probably because it is followed by a local ~24% SD in the next 25-nt that could act as a strong heterologous barrier. The other endpoints were located in regions below MEPS which are usually followed by barriers with a higher SD (*e.g*., endpoint at position 1371 is preceded by 25-nt with ~24% SD, at position 1251 by ~24% SD). No patched sequences were observed, discarding that in some transformants multiple recombination events had occurred at different loci. As in *rec^+^*, the recombination endpoints in the *recF*15, Δ*recO*, Δ*recU*, Δ*ruvAB* and Δ*addAB* transformants did not always coincide with the longest MEPS present (Table S1). In these mutants one endpoint was usually at a region where a MEPS was located, whereas the other endpoint was less specific, and often below MEPS. All these results suggest that a MEPS is necessary to initiate strand invasion, but a SD barrier may halt RecA-mediated uni- or bidirectional DNA strand exchange.

**Figure 4.**
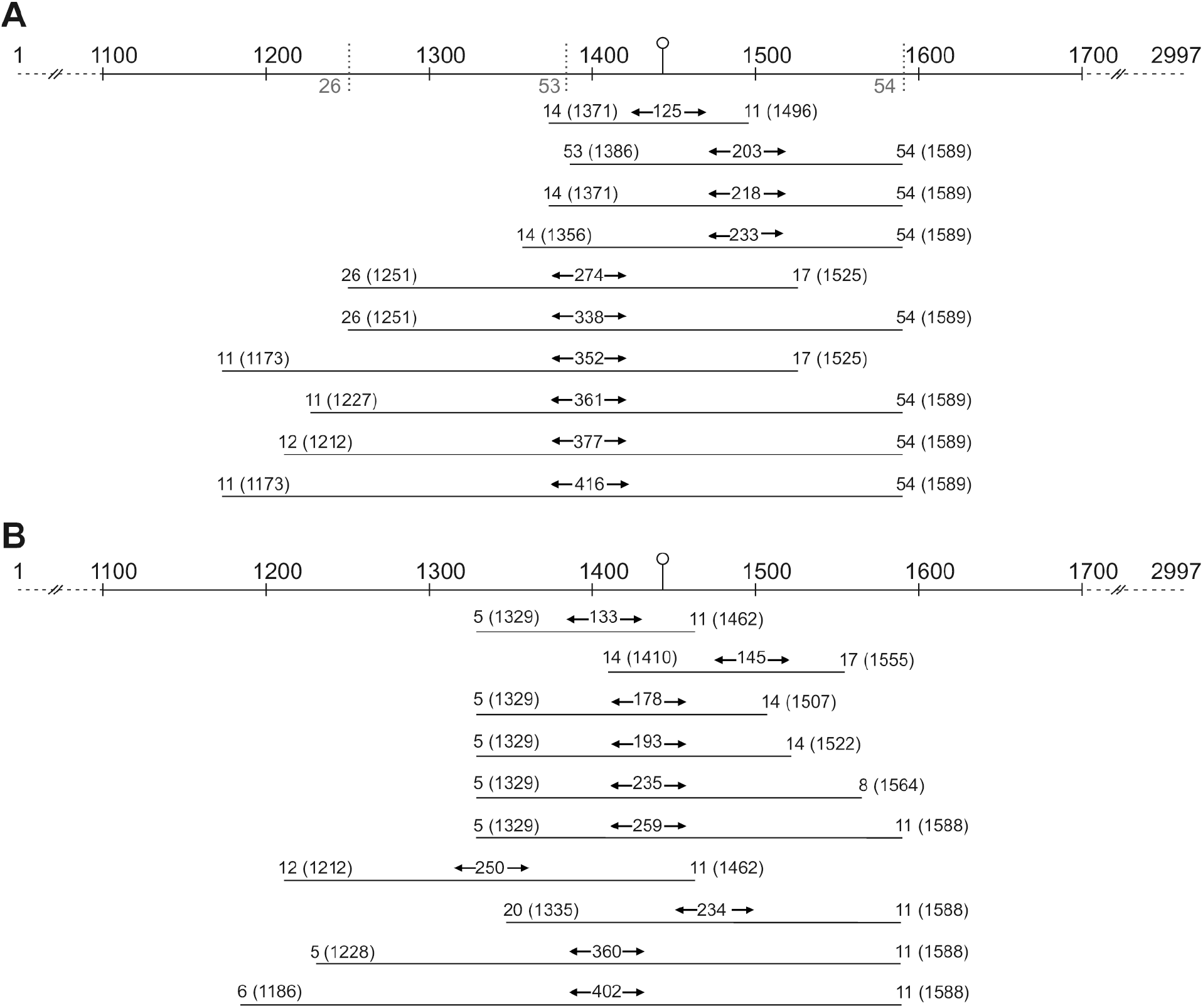
Mapping of recombination endpoints during interspecies CT in *rec^+^* cells. The length of integrated DNA and recombination endpoints were determined by sequencing ten different R^R^ clones obtained following transformation with *Bam* DSM7 *rpoB482* donor DNA (10.12% SD, in A) or *Bli* DSM13 *rpoB*482 donor DNA (14.52% SD, in B). In the top of the figure, the features of the donor DNA are indicated: The mutation that confers Rif^R^, located at 1443 bp in the *rpoB* genes, is marked by a pin. In *Bam* DSM7 *rpoB*482 DNA (10.12% SD) the regions in the sequence where MEPSs (i.e., fully homologous regions larger than 26-bp) are found are denoted by a vertical dotted line. Their length is indicated also. MEPS regions are not observed in *Bli* DSM13 *rpoB482* DNA (14.52 % SD). The MEPS and integration lengths of ten different Rif^R^ clones are shown, in brackets are indicated the location of recombination endpoints.

Nucleotide sequence analysis of 15-30 Rif^R^ clones obtained with ~15% SD in *ΔruvAB* and *rec^+^* cells showed that all were genuine transformants, but these values dropped to ~40-50% in Δ*recU*, Δ*recO*, Δ*addAB*, and *recF*15. The mean integration length was between 400- to 130-nt in *recF*15, Δ*recO*, Δ*recU*, Δ*ruvAB* and *ΔaddAB*, as well as in the *rec^+^* control (Table S1 and Fig. 4B). This donor DNA has a MEPS of 104-nt (position 54-157) upstream of the *rpoB*482 mutation, and a 38-nt MEPS (position 2295-2332) downstream, but *~*2100-nt inserts were not observed, confirming that the region between the MEPS are relevant. Potential MEPS are not present in the integrated region (Fig. 4B). The sequence analysis of *rec^+^* transformants revealed a great heterogenicity in the recombination endpoints, as well as in the length of the MEPS. In all cases the MEPS used were short: between 5- to 20-nt without mismatches (Fig. 4B and Table S1). However, if 1-mismatch is allowed a longer MEPS is detected (*e.g*., the endpoint at position 1329 increases from 5- to 26-nt) (Fig. 4B).

Beyond 15% SD the transformation efficiency and the frequency of spontaneous mutation almost overlap, although there is a ~3-fold difference, and it is higher than in a *ΔrecA* mutant, suggesting that still this is a RecA-dependent integration event (Fig. 1). Four regions at or above MEPS (74-, 56-, 28- and 26-nt) upstream and one downstream (26-nt) of the *rpoB*482 mutation are present in the 17% SD donor, but they were not used to integrate the region in-between.

Nucleotide sequence analyses of Rif^R^ *rec^+^* clones obtained with ~17% SD showed that only ~37% were genuine transformants, *i.e*, they had incorporated two or more nucleotides of donor DNA. Similarly, 20-30% of the Rif^R^ clones obtained in *ΔrecO, ΔaddAB, ΔrecU, ΔruvAB* and *recF15* were genuine transformants (Fig. 3A-B), and the mean integration length was also 4- to 8-nt, or ~5-fold below MEPS (Table S1). The observed low efficiency of micro-homologous integration cannot be attributed to a defect in the resulting RpoB482 protein, because at *~*17% SD, except the amino acid change that confers Rif^R^, there is no mutation in the 50 residues intervals up and downstream the Rif^R^ change (Fig. S1C). The close inspection in donor DNA with 17% SD of the region surrounding the *rpoB*482 mutation showed that this mutation is embedded in a region with strong SD with recipient DNA. The 25-nt region upstream of the *rpoB*482 mutation has 8 mismatches (*i*.*e*, 32% SD) and the one downstream 10 mismatches (40% SD) that could act as heterologous barriers.

At ~21% SD there are two sites at or above MEPS (42-nt [at position 90-131] and 27-nt [at position 2781-2807]), one on each side of the *rpoB*482 mutation, but they were not used. After sequencing of transformants obtained in *recF*15, Δ*recO*, Δ*recU*, Δ*ruvAB*, Δ*addAB* or *rec*^+^, we found that again just few nucleotides from the donor had been integrated, and that the fraction of genuine transformants was ~20% (Fig 3A-B). Here, the analysis showed that the *rpoB*482 mutation is also surrounded by higher SD: 8 mismatches in 25-nt upstream (*i*.*e*, ~32% SD) and the one downstream 6 mismatches (*i*.*e*, ~24% SD). Finally, at ~23% SD, only one MEPS exists (at position 801-826). The proportion of genuine Rif^R^ transformants accounts to only ~6% of the sequenced clones in the *rec*^+^ strain. The mean integration length in *rec*^+^ transformants was 4- to 8-nt (Fig. 3A-B). Similar results were obtained in *recF*15, Δ*recO*, Δ*recU*, Δ*ruvAB* and Δ*addAB*.

### RecD2, RecJ, RecX, RadA/Sms and DprA are crucial for interspecies CT

Nucleotide sequence analyses of 10-30 Rif^R^ clones obtained in the Δ*radA*, Δ*recJ*, Δ*recX*, Δ*recD2* or Δ*dprA* context revealed that when divergence was low (*~*2%), the mean integration length of the *rpoB*482 DNA was similar to the *rec^+^* control (Fig 3C-D). The proportion of genuine transformants was different in these mutants. At ~8% SD, all sequenced clones obtained in *DradA* were genuine transformants, but this number was reduced to ~50 % in Δ*dprA*, Δ*recJ* and Δ*recX*, and even lowered to ~25% in Δ*recD2*. Representatives of the genuine Rif^R^ clones are documented in Table S2.

The efficiency of interspecies transformation at ~10% SD was strongly reduced in these mutants, to levels similar to the spontaneous mutation rate (Fig. 1). Only ~25% of the Rif^R^ clones obtained in Δ*recX* were genuine transformants, and these values lowered to ~15% in Δ*recD2*, and ~5% in Δ*recJ*and Δ*radA* cells. Furthermore, in the Δ*dprA* strain all 25 sequenced Rif^R^ clones were spontaneous mutants (Table S2). The mapping of recombination endpoints in the Δ*recJ*, Δ*recD2*, Δ*recX* and Δ*radA* transformants showed that, as in the *rec^+^* transformants, the 3’-endpoint of recombination was located at the 54-nt MEPS, except in some Δ*recX* transformants. The 5’-endpoint of recombination was usually located at a larger MEPS when compared with the *rec^+^* control (Tables S1 and S2).

At ~15% SD, all sequenced Rif^R^ clones obtained were spontaneous mutants in the Δ*recX*, Δ*radA*, Δ*recJ* and Δ*dprA* strains. In Δ*recD2*, only one of the sequenced Rif^R^ clones (1/19) was a genuine transformant with a mean integration length >120-nt (Table S2). At ~17% SD, all sequenced Rif^R^ clones were spontaneous mutants in the Δ*recX*, Δ*radA*, Δ*recJ*, Δ*dprA*, and Δ*recD2* backgrounds. These results show that these functions are all needed for homology-directed integration when SD is larger than 10%, and also for micro-homologous integration.

## Discussion

From the data presented in this work, it can be inferred that recombination proteins differently facilitate adaptation and genetic diversity, and that two different recombination mechanisms occur depending on the SD of the interspecies DNA. Up to 15% SD, homology-directed HR accounts for the integration of divergent sequences longer than 130-nt. Beyond this, integration of just few nucleotides at micro-homologous segments is observed.

Inactivation of the main end-resection complex (*addAB*), a positive mediator (*recO*), a positive (*recF*) or a negative (*recU*) modulator or enzymes essential for Holliday junction processing and cleavage (*ruvAB*, *recU*) barely affects heterogamic CT (Fig. 1A-B), although they are essential for DNA DSB repair ^39,43,44^. The interpretation of such result may not be so simple, because cells lacking both RecF and AddAB are blocked even in homogamic CT ^43^, suggesting that they are backup functions during CT (*e.g*., RecF is essential in the *recX* context) ^25^. We have observed that a specific subset of HR proteins (RecA, DprA, RecX, RadA/Sms, RecD2 and RecJ) contributes to acquire homeologous DNA. Except DprA which is competence specific and present in all transformable bacteria, and even in non-transformable ones ^10^, the recombination proteins identified here also participate, together with another subset of HR proteins, in the accurate repair of lethal DSBs and in the restart of stalled replication forks. In some distantly related competent cells (*e.g*., Proteobacteria phylum) RecD2 is absent, and RadA/Sms is replaced by ComM ^45^. They differently participate in the recombination mechanisms that may be active during the acquisition of homeologous DNA. In the absence of RecA CT is blocked ^16^. Interspecies homology-directed CT was blocked with DNA of the same clade (up to ~8% SD) in Δ*dprA*, beyond ~10% SD in the *ΔrecX, ΔrecJ* and Δ*radA* backgrounds, and beyond ~15% SD in Δ*recD2* cells (Fig. 3C-D), showing an essential role for these proteins in the acquisition of interspecies DNA.

We can envision that during homology-directed CT, RecA nucleates in the incoming ssDNA coated by SsbA and SsbB, with the help of the DprA-SsbA two-component mediator. Then, a dynamic RecA nucleoprotein filament, with the contribution of DprA-SsbA and RecX, identifies an identical sequence in the recipient genome. Once a homologous region is found RecA promotes DNA strand invasion to produce a metastable heteroduplex DNA ^23,27^. RecA at this D-loop interacts with and loads the branch-migration translocase RadA/Sms as documented in Firmicutes ^34,46^. RadA/Sms, in concert with RecA, facilitates bidirectional branch migration until a region with a SD >20% is found. This is consistent with the *in vitro* observation that homology-directed RecA-mediated strand-exchange halted at DNA patches >16% SD ^16^.

The contribution of RecD2 helicase and RecJ exonuclease to homology-directed recombination during interspecies CT is poorly understood. Recently, it was suggested that RecD2 contributes to branch migrate the heteroduplex DNA in a relaxed molecule, perhaps upon cleavage of the displaced strand ^28^. As RecJ of other naturally competent cells ^47^, *B. subtilis* RecJ possesses an extra C-terminal domain, absent in *E. coli* RecJ, critical for protein-protein interaction (*e.g*., SsbA). Perhaps in concert with the RecD2 helicase, it might degrade the displaced strand and the non-paired tails. Finally, the ends are sealed, leading to the acquisition of homeologous DNA >130-nt.

It was remarkable the heterogeneity of the MEPS at recombination endpoints in homology-directed HR. *In vitro* assays with *E. coli* RecA show that identity search relies on probing tracts of 8-nt homology, based on the transient interactions between the stretched ssDNA within the filament and bases in a locally melted and stretched DNA duplex ^48,49^. It has been shown that RecA evolved to tolerate 1-nt mismatch every ~8-nt region *in vitro*, albeit DNA strand exchange with a short region of 16% SD is delayed ^16,50^. We propose that a sum of delays might compromise DNA strand exchange and determines the length of DNA integrated.

Beyond 15% sequence, integration of <10-nt segments was observed in Δ*addAB*, *recF*, Δ*recO*, Δ*recU* and Δ*ruvAB* (Fig. 3A-B), but it was not detected in Δ*recA*, Δ*dprA*, Δ*recJ*, Δ*recX*, Δ*radA* and Δ*recD2* cells (Fig. 3C-D). These results showed that a similar set of recombination functions are also required for this short integration, which occurs up to 23% SD, although with low efficiency. Here we propose a homology-facilitated micro-homologous integration mechanism: As above, a RecA dynamic filament, with the help of DprA-SsbA and RecX, searches for and identifies a MEPS on the *rpoB*482 DNA, which is used in this case as an anchor region, to produce a metastable D-loop intermediate, in concert with RadA/Sms. Once the DNA is anchored at MEPS, DprA could mediate the annealing of short stretches of homeologous DNA (3- to 8-nt) around the *rpoB*482 mutation ^23^. Then, the donor ssDNA loop between the anchored region and the micro-homologous paired segment has to deleted, perhaps by RecJ in concert with RecD2. Finally, the ends of the integrated segment are sealed and rapidly expressed ^45^. This mechanism differs from HFIR observed in replicating competent *S. pneumoniae* and *A. baylyi* cells (see Introduction) ^20,21^. In these competent bacteria, inactivation of *recBCD* or *recJ* significantly increased HFIR ^19,21^. In contrast, in non-replicating competent *B. subtlis* cells integration of thousands of nucleotides of the heterogamic DNA with the subsequent deletion of the recipient DNA was not observed, and inactivation of *recJ* blocked homology-facilitated micro-homologous integration. Competent *A. baylyi* cells can also integrate short ssDNA (20-nt) in the recipient genome by another mechanism, which occurs with extremely low efficiency.

This mechanism requires active DNA replication, inactivation of *recJ* and is independent of RecA ^51^. It is likely that competent *B. subtilis* cells may use homology-facilitated micro-homologous integration to restore genes inactivated by mutations and thereby prevent the irreversible deterioration of genomes (Muller’s ratchet) ^4^. At ~23% SD interspecies CT frequency was similar to spontaneous mutations, suggesting that beyond *~*23% SD micro-homologous integration might be inefficient, probably because the unique MEPS present is too short to serve as a stable anchor region ^19^. Similarly, competent *H. pylori* cells cannot be transformed by *Campylobacter jejuni* DNA with ~24% SD ^52^.

Can the above observations be extrapolated to interspecies Hfr chromosomal conjugation? The frequency of interspecies Hfr chromosomal conjugation also decreases log-linearly with increased SD up to ~16% SD. Inactivation of *mutSL* alleviates interspecies Hfr conjugation by ~1000-fold, and deletion of *recBCD* or *ruvAB* reduced interspecies Hfr conjugation, but inactivation of *recJ* increases it ^2,3,53,54^, suggesting that interspecies Hfr chromosomal conjugation uses the repair-by-recombination mechanism. In contrast, inactivation of *mutSL* marginally prevents interspecies CT with up to 15% SD in Firmicutes ^15,16,55,56^ and inactivation of *recJ*, but not *addAB* or *ruvAB*, inhibits interspecies CT. All these results suggest that bacteria have evolved different genetic recombination mechanisms devoted to interspecies genetic exchange to generate diversity. Bacterial *recA*, *radA* and *recX* genes, which play crucial roles in interspecies CT, perhaps contributed in their transfer from mitochondria or chloroplasts to the nucleus of land plants, green algae and moss ^57^, although the evolutionary force and molecular functions that contributed to the transfer of these genes well beyond the species boundaries is poorly understood.

## Materials and methods

### Bacterial strains and donor DNAs

The parental strain was *B. subtilis* BG1359. The *rec* mutations listed in Table 1 were introduced by SPP1 transduction ^58^.

The *rpoB* gene from different species, which encodes for the ²-subunit of RNA polymerase, was used as donor DNA, and the *rpoB*482 mutation, which renders cells Rif^R^, was introduced into all the donor DNAs (*Annex* 2, supplementary material). Plasmid DNA was prepared by Qiagen extraction and extensive dialysis in Tris-EDTA buffer ^17^.

### Transformation assays

Natural competence was induced as described ^42^. Competent cells were incubated with 0.1 μg·ml^-1^ of the indicated *rpoB*482 donor DNA (30 min, 37°C), and then plated on Rif (8 μg·ml^-1^) containing LB-agar plates. A control was performed in which competent cells were treated equally, but with no donor DNA to score the appearance of spontaneous Rif^R^ mutants. These values (spontaneous Rif^R^ mutants) were extracted to the number of transformants. Transformation frequency was calculated as the number of Rif^R^ transformants per colony-forming-unit (CFU).

### Mapping of integration endpoints

The integration endpoint is defined by the end of the donor sequence followed by the sequence of the recipient. To map integration endpoints, the *rpoB* gene from the Rif^R^ transformants was amplified by PCR, and its nucleotide sequence compared with the one of recipient and donor strains. The presence or the absence of the mismatches between the donor and the recipient DNA were used to determine the MEPS. Endpoints are defined as described ^59^, and integration length is calculated as the distance between endpoints ^59^.

## Acknowledgements

ES and CR thank the Ministerio de Ciencia e Innovación [Agencia Estatal de Investigación] (MCIU[AEI]), BES-2013-063433 to E.S., and BES-2017-080504 to C.R. for the fellowships. This work was partially supported by MCIU/AEI/FEDER, EU PGC2018-097054-B-I00 to J.C.A and S.A. The funders had no role in study design, data collection and analysis, decision to publish, or preparation of the manuscript.

## Author contributions

S.A. and J.C.A. conceived the project and designed the experiments, E.S., C.R. performed the experiments, E.S., C.R. S.A. and J.C.A. evaluated the results, S.A. and J.C.A. wrote the manuscript.

