## Supplementary information for "Recombination proteins differently control the acquisition of homeologous DNA during *Bacillus subtilis* natural chromosomal transformation"

This supplementary material contains

- Supplementary Annexes (1 to 3)
- Supplementary references
- Supplementary Tables (S1 and S2)
- Supplementary Figure S1

[1]Department of Microbial Biotechnology, Centro Nacional de Biotecnología, CNB-CSIC, 3 Darwin Str, 28049 Madrid, Spain.
✉ Juan C. Alonso and Silvia Ayora

**Annex 1**. Why *Bacillus subtilis* was selected as the experimental system

*B. subtilis* competent cells selected to analyse how the recombination machinery limits heterogamic chromosomal transformation (CT): first, its uptake apparatus internalises any environmental DNA with similar efficiency [1,2]. Second, the recombination between the incoming ssDNA and the non-replicating haploid genome of a competent cell simplifies the interpretation of the data [1,3]. Finally, these cells show no codon usage preferences [4].

**Annex 2**. Donor DNAs

In all cases, a 2997-bp fragment of the *rpoB*482 gene was *in vitro* synthesized and cloned into the *E. coli* pUC57 plasmid [5]. The *rpoB*482 mutation is a single C to T transition, that confers Rif$^R$, but does not affect RNA polymerase activity. For homogamic CT, plasmid pCB980-borne *rpoB*482 of *B. subtilis* 168 (*Bsu* 168 [1 mismatch]) was used. For heterogamic CT the 2997-bp *rpoB*482 donor DNA was derived from: i) bacteria of the *B. subtilis* clade, with up to ~8% sequence divergence (SD) (pCB981-borne *rpoB*482 of *B. subtilis* W23 (*Bsu* W23 [2.47% SD, 74 mismatches]), pCB982-borne *rpoB*482 of *B. atrophaeus* 1942 (*Bat* 1942, 8.35% SD, 250 mismatches]), ii) a bacterium of the *B. amyloliquefaciens* clade with 10.12% SD (pCB983-borne *rpoB*482 of *B. amyloliquefaciens* DSM7 (*Bam* DSM7, 303 mismatches]); iii) bacteria of the *B. licheniformis* clade: with 14.52% (pCB984-borne *rpoB*482 of *B. licheniformis* DSM13 [*Bli* DSM13, 435 mismatches]) and 17% SD (pCB1054-borne *rpoB*482 of *B. gobiensis* FJAT4402 (*Bgo* FJAT4402, 510 mismatches/insertion/deletion]); iv) a bacterium of the *B. thuringiensis* clade with 20.83% SD (pCB985-borne *rpoB*482 of *B. thuringiensis* MC28 [*Bth* MC28, 624 mismatches/insertion/deletion]); and v) a far distant *Bacillus* with 22.74% SD (pCB1056-borne *rpoB*482 DNA of *B. smithii* DSM4216 [*Bsm* DSM4216, 681 mismatches/insertion/deletion]): All donor *rpoB*482 DNAs have similar dG + dC content. Plasmids were purified and used as donor DNA. This *E. coli* plasmid cannot replicate into *B. subtilis* cells, but the homologous *rpoB*482 DNA integrates and confers Rif$^R$ upon CT.

Previously, it has been shown that expression of a plasmid-borne *B. thuringiensis rpoB*482 gene, which confers Rif$^R$ and express a RpoB protein with 20.83% SD, shows no apparent fitness cost in the recipient strain [5]

**Annex 3**. RecD2 controls the appearance of spontaneous mutations

The rate of appearance of spontaneous Rif$^R$ mutations was assessed in the different strains listed in Table 1. In the different *rec*$^-$ strains, the number of Rif$^R$ colonies that appeared in the absence of *rpoB*482 DNA was similar to the *wt* control, except in Δ*recD2* cells. The mutation frequencies were similar in the Δ*rok* or *rok*$^+$ backgrounds (~7 to 9 x $10^{-9}$). However, the frequency of appearance of Rif$^R$ colonies increased ~3-fold (~2 to 4 x $10^{-8}$) in competent Δ*rok* Δ*recD2*, as in Δ*recD2* cells [6]. Similarly, inactivation of *recD*2 increased 3.6-fold the frequency of spontaneous trimethoprim resistant mutants, and the mutation frequency was also increased in *B. anthracis recD2* mutants [7,8]. It becomes clear that RecD2 controls the appearance of spontaneous mutations by an unknown mechanism.

**Table S1.** Mean integration length during interspecies chromosomal transformation in *ΔaddAB, ΔrecO, recF15, ΔruvAB* and *ΔrecU* mutants

| Genetic background | Divergence (in %) | Left end (MEPS in bp) | Right end (MEPS in bp) | Integration (in bp)[a] |
|---|---|---|---|---|
| *ΔaddAB* | 8.35 | 792 (50) | 1647 (17) | 855 |
| | | 780 (11) | 1589 (81) | 809 |
| | | 1128 (23) | 1867 (41) | 739 |
| | | 1152 (23) | 1729 (23) | 577 |
| | | 1407 (36) | 1867 (41) | 460 |
| | 10.12 | 1128 (23) | 1771 (26) | 643 |
| | | 1152 (20) | 1771 (26) | 619 |
| | | 1251 (26) | 1525 (17) | 274 |
| | | 1356 (14) | 1589 (54) | 233 |
| | | 1371 (14) | 1589 (54) | 218 |
| | 14.52 | 1335 (20) | 1588 (11) | 253 |
| | | 1335 (20) | 1564 (8) | 229 |
| | | 1335 (20) | 1555 (17) | 220 |
| | | 1335 (20) | 1522 (14) | 187 |
| | | 1410 (14) | 1588 (11) | 178 |
| | | - | - | (5)[a] |
| | 17.0 | 1440 | 1448 | ~8 |
| | | 1441 | 1445 | ~4 |
| | | 1441 | 1445 | ~4 |
| | | 1441 | 1445 | ~4 |
| | | - | - | (3)[a] |
| *ΔrecO* | 8.35 | 1251 (18) | 2132 (33) | 881 |
| | | 780 (11) | 1589 (81) | 809 |
| | | 1128 (23) | 1867 (41) | 739 |
| | | 1128 (23) | 1729 (23) | 601 |
| | | 1128 (23) | 1647 (17) | 519 |
| | 10.12 | 1117 (12) | 1589 (54) | 472 |
| | | 1300 (17) | 1663 (11) | 363 |
| | | 1300 (17) | 1589 (54) | 289 |
| | | 1356 (14) | 1589 (54) | 233 |
| | | 1371 (14) | 1589 (54) | 218 |
| | 14.52 | 1309 (11) | 1555 (17) | 246 |
| | | 1335 (20) | 1555 (17) | 220 |
| | | 1335 (20) | 1522 (14) | 187 |
| | | 1410 (14) | 1588 (11) | 178 |
| | | 1410 (14) | 1555 (17) | 145 |
| | | - | - | (4)[a] |
| | 17.0 | 1441 | 1448 | ~7 |
| | | 1441 | 1445 | ~4 |
| | | 1441 | 1445 | ~4 |
| | | 1441 | 1445 | ~4 |
| | | - | - | (3)[a] |
| | 8.35 | 1251 (18) | 2132 (33) | 881 |
| | | 780 (11) | 1589 (81) | 809 |
| | | 792 (50) | 1589 (81) | 797 |
| | | 954 (35) | 1729 (23) | 775 |
| | | 954 (35) | 1589 (81) | 635 |
| | 10.12 | 1128 (23) | 1771 (26) | 643 |
| | | 1250 (26) | 1615 (11) | 365 |
| | | 1251 (26) | 1589 (54) | 339 |
| | | 1371 (14) | 1663 (11) | 292 |

| | | | | |
|---|---|---|---|---|
| *recF15* | | 1371 (14) | 1589 (54) | 218 |
| | 14.52 | 1212 (12) | 1690 (14) | 478 |
| | | 1251 (14) | 1715 (11) | 464 |
| | | 1374 (8) | 1690 (14) | 316 |
| | | 1386 (8) | 1588 (11) | 202 |
| | | 1335 (20) | 1522 (14) | 187 |
| | | - | - | (5)[a] |
| | 17.0 | 1441 | 1448 | ~7 |
| | | 1441 | 1445 | ~4 |
| | | 1441 | 1445 | ~4 |
| | | 1441 | 1445 | ~4 |
| | | - | - | (2)[a] |
| *ΔruvAB* | 8.35 | 1342 (11) | 1589 (81) | 211 |
| | | 954 (35) | 1589 (81) | 635 |
| | | 1128 (23) | 1729 (23) | 601 |
| | | 1152 (23) | 1729 (23) | 577 |
| | | 1407 (36) | 1867 (41) | 460 |
| | 10.12 | 1128 (23) | 1771 (26) | 643 |
| | | 1150 (20) | 1771 (26) | 621 |
| | | 1250 (26) | 1525 (17) | 275 |
| | | 1371 (14) | 1615 (11) | 244 |
| | | 1371 (14) | 1589 (54) | 218 |
| | 14.52 | 1335 (20) | 1690 (14) | 355 |
| | | 1335 (20) | 1588 (11) | 253 |
| | | 1335 (20) | 1555 (17) | 220 |
| | | 1335 (20) | 1507 (14) | 172 |
| | | 1410 (14) | 1555 (17) | 145 |
| | 17.0 | 1439 | 1445 | ~6 |
| | | 1441 | 1448 | ~7 |
| | | 1441 | 1445 | ~4 |
| | | 1441 | 1445 | ~4 |
| | | - | - | (5)[a] |
| *ΔrecU* | 8.35 | 780 (11) | 1589 (81) | 809 |
| | | 1128 (23) | 1867 (41) | 739 |
| | | 954 (35) | 1589 (81) | 635 |
| | | 1128 (23) | 1729 (23) | 601 |
| | | 1407 (36) | 1867 (41) | 460 |
| | 10.12 | 1128 (23) | 1771 (26) | 643 |
| | | 1128 (23) | 1589 (54) | 461 |
| | | 1250 (26) | 1525 (17) | 275 |
| | | 1334 (11) | 1589 (54) | 255 |
| | | 1356 (14) | 1589 (54) | 233 |
| | 14.52 | 1335 (20) | 1690 (14) | 355 |
| | | 1335 (20) | 1648 (11) | 313 |
| | | 1335 (20) | 1555 (17) | 220 |
| | | 1335 (20) | 1522 (14) | 187 |
| | | 1410 (14) | 1555 (17) | 145 |
| | | - | - | (3)[a] |
| | 17.0 | 1441 | 1448 | ~8 |
| | | 1441 | 1445 | ~4 |
| | | 1441 | 1445 | ~4 |
| | | 1441 | 1445 | ~4 |
| | | - | - | (3)[a] |

Five Rif$^R$ clones for each condition are documented. [a]The number of spontaneous Rif$^R$ clones analysed (non-genuine transformants) are denoted between parentheses.

**Table S2.** Mean integration length during interspecies chromosomal transformation in *ΔrecD2*, *ΔrecX*, *ΔradA ΔrecJ* and *ΔdprA* mutants

| Genetic background | Divergence (in %) | Left end (MEPS in bp) | Right end (MEPS in bp) | Integration (in bp) [a] |
|---|---|---|---|---|
| *ΔrecD2* | 8.35 | 633 (11) | 1969 (20) | 1336 |
| | | 1128 (23) | 1589 (81) | 461 |
| | | 1394 (11) | 2980 (13) | 1585 |
| | | 1370 (36) | 1589 (81) | 219 |
| | | - | - | (13)[a] |
| | 10.12 | 805 (29) | 1589 (54) | 784 |
| | | 1151 (20) | 1589 (54) | 438 |
| | | 1371 (14) | 1589 (54) | 218 |
| | | 1386 (53) | 1589 (54) | 203 |
| | | - | - | (21)[a] |
| | | 1395 (11) | 1555 (17) | 160 |
| | 14.52 | - | - | (18)[a] |
| | 17.0 | - | - | (25)[a] |
| *ΔrecX* | 8.35 | 1152 (23) | 1729 (23) | 574 |
| | | 1116 (12) | 1589 (81) | 473 |
| | | 1407 (36) | 1867 (41) | 460 |
| | | 1344 (20) | 1589 (81) | 245 |
| | | - | - | (5)[a] |
| | 10.12 | 1251 (26) | 1589 (54) | 338 |
| | | 1251 (26) | 1525 (17) | 274 |
| | | 1335 (20) | 1615 (11) | 280 |
| | | 1371 (14) | 1525 (17) | 154 |
| | | - | - | (12)[a] |
| | 14.52 | - | - | (25)[a] |
| *ΔradA* | 8.35 | 1 (158) | 1909 (23) | 1909 |
| | | 780 (11) | 1589 (81) | 809 |
| | | 792 (50) | 1589 (81) | 797 |
| | | 954 (35) | 1729 (23) | 775 |
| | | 954 (35) | 1589 (81) | 635 |
| | 10.12 | 1371 (14) | 1589 (54) | 218 |
| | | - | - | (22)[a] |
| | 14.52 | - | - | (30)[a] |
| *ΔrecJ* | 8.35 | 407 (77) | 1774 (11) | 1367 |
| | | 1152 (23) | 1589 (81) | 437 |
| | | 1370 (23) | 1589 (81) | 219 |
| | | 1370 (23) | 1729 (23) | 359 |
| | | 1370 (23) | 1639 (17) | 269 |
| | | - | - | (5)[a] |
| | 10.12 | 1438 (53) | 1589 (54) | 151 |
| | | - | - | (20)[a] |
| | 14.52 | - | - | (25)[a] |
| *ΔdprA* | 8.35 | 1128 (23) | 1867 (41) | 739 |
| | | 954 (35) | 1589 (81) | 635 |
| | | 1251 (14) | 1774 (11) | 523 |
| | | 1407 (36) | 1867 (41) | 460 |
| | | - | - | (6)[a] |
| | 10.12 | - | - | (25)[a] |
| | 14.52 | - | - | (25)[a] |
| | 17.0 | - | - | (20)[a] |

When available 5 $Rif^R$ clones for each condition are documented. [a]The number of spontaneous $Rif^R$ clones analysed (non-genuine transformants) are denoted between parentheses.

**Figure S1**

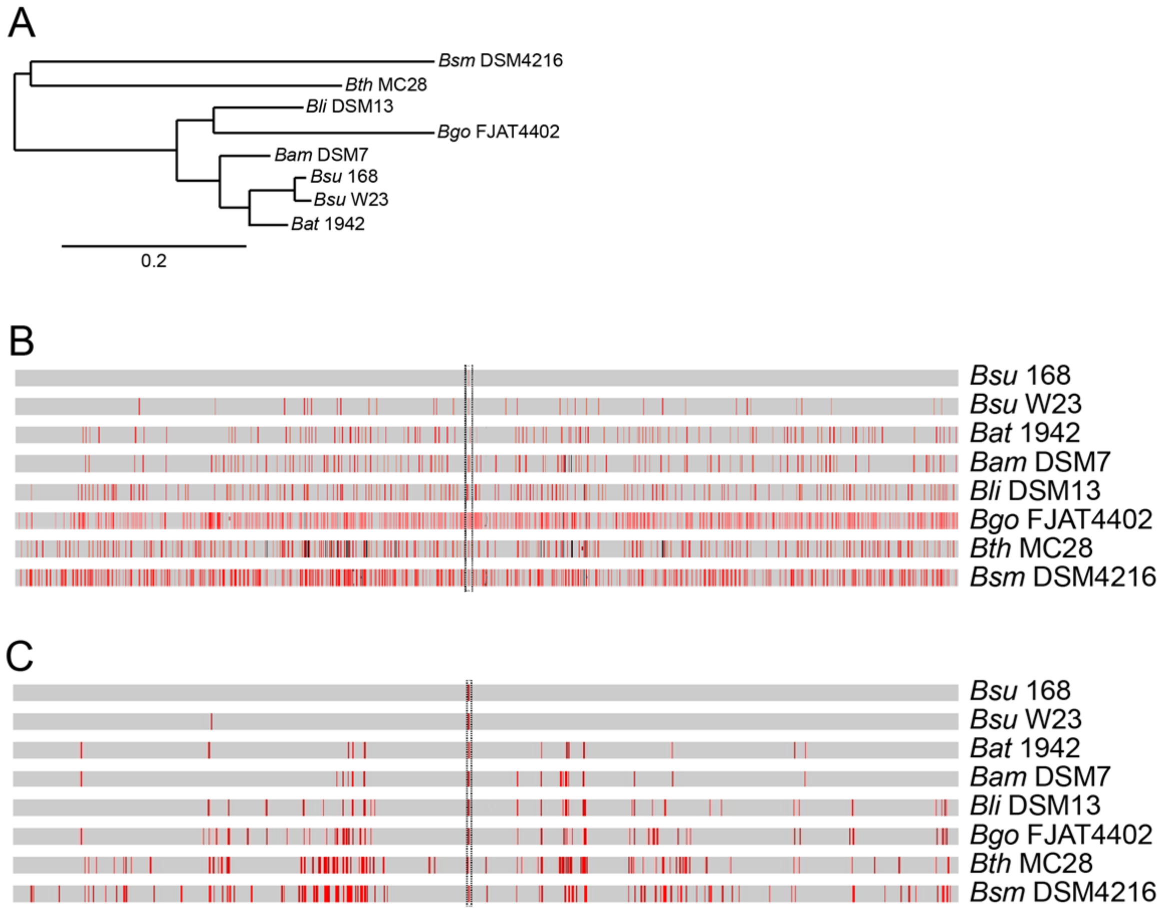


**Supplementary Figure 1**. Distribution of sequence divergence among different *Bacillus* species. (A) Phylogenetic tree of the selected *Bacillus* species or subspecies based on the *rpoB* nucleotide sequence. Branching confidence values are based on 1,000 bootstrap replicates. (B) Nucleotide sequence differences between the different *Bacilli* genes. A single C-to-T transition mutation at codon 482 (framed by dotted lines), which is centrally located (at position 1443) in the *rpoB*482 gene confers *Rif*$^{R}$. The *rpoB*482 DNA was derived from *B. subtilis* 168 (*Bsu* 168), *B. subtilis* W23 (*Bsu* W23), *B. atrophaeus* 1942 (*Bat* 1942), *B. amyloliquefaciens* DSM7 (*Bam* DSM7), *B. licheniformis* DSM13 (*Bli* DSM13), *B. gobiensis* FJAT-4402 (*Bgo* FJAT4402), *B. thuringiensis* MC28 (*Bth* MC28) and *B. smithii* DSM4512 (*Bsm* DSM4216). Mismatches between donor and recipient are indicated by vertical red bars, and insertions/deletions by vertical black bars. Bar thickness represents the number of mismatches in the neighbourhood. (C) Sequence alignment of the RpoB protein from *Bsu* 168, *Bsu* W23, *Bat* 1942, *Bam* DSM7, *Bli* DSM13, *Bgo* FJAT4402, *Bth* MC28 and *Bsm* DSM4216. The proteins have a high degree of sequence identity (grey). Conserved replacements are indicated by vertical red bars and non-conserved residues by black bars. The alignments were performed using BLAST.
